# Single cell transcriptomic analysis of renal allograft rejection reveals novel insights into intragraft TCR clonality

**DOI:** 10.1101/2023.02.08.524808

**Authors:** Tiffany Shi, Ashley R. Burg, J. Timothy Caldwell, Krishna Roskin, Cyd M. Castro-Rojas, P. Chukwunalu Chukwuma, George I. Gray, Sara G. Foote, Jesus Alonso, Carla M. Cuda, David A. Allman, James S. Rush, Catherine H. Regnier, Grazyna Wieczorek, Rita R. Alloway, Adele R. Shields, Brian M. Baker, E. Steve Woodle, David A. Hildeman

## Abstract

Bulk analysis of renal allograft biopsies (rBx) identified RNA transcripts associated with acute cellular rejection (ACR); however, these lacked cellular context critical to mechanistic understanding. We performed combined single cell RNA transcriptomic and TCRα/β sequencing on rBx from patients with ACR under differing immunosuppression (IS): tacrolimus, iscalimab, and belatacept. TCR analysis revealed a highly restricted CD8^+^ T cell clonal expansion (CD8_EXP_), independent of HLA mismatch or IS type. Subcloning of TCRα/β cDNAs from CD8_EXP_ into Jurkat76 cells (TCR^-/-^) conferred alloreactivity by mixed lymphocyte reaction. scRNAseq analysis of CD8_EXP_ revealed effector, memory, and exhausted phenotypes that were influenced by IS type. Successful anti-rejection treatment decreased, but did not eliminate, CD8_EXP_, while CD8_EXP_ were maintained during treatment-refractory rejection. Finally, most rBx-derived CD8_EXP_ were also observed in matching urine samples. Overall, our data define the clonal CD8^+^ T cell response to ACR, providing novel insights to improve detection, assessment, and treatment of rejection.

## Introduction

Transplantation is the most effective treatment for kidney failure, providing improved survival and quality of life. To prevent rejection, patients are maintained on lifelong immunosuppression (IS) therapy, the standard-of-care being the calcineurin inhibitor (CNI) tacrolimus. Although current CNI-based regimens provide low acute rejection rates (~10%) in the first post-transplant year, thereafter patients remain at risk of therapeutically resistant late rejection that occurs at a rate of 1-3% per year. In addition, CNI-based regimens are associated with substantial toxicities due to off-target effects^1,2^. To avoid CNI toxicities, targeted IS agents that block T cell co-stimulation are being developed. Belatacept (CTLA4-Ig) is FDA-approved, and despite increased rejection rates 1-year post-transplant, long-term kidney function is increased under belatacept IS^3^. However, belatacept-refractory rejection (BRR) episodes are more severe and more difficult to treat than rejection episodes that occur under CNI-based IS^4-6^. Antibody-mediated blockade of the CD40/CD40L co-stimulatory pathway has shown promise as maintenance IS in non-human primates^7-9^ and in pig-to-monkey xenografts^10-12^. Fully human anti-CD40 monoclonal antibodies, such as iscalimab and bleselumab, have provided effective IS in human trials, but remained less efficacious compared to tacrolimus-based treatment in the prevention of organ rejection^13,14^. Thus, although co-stimulatory blockade approaches are promising, less toxic IS regimens, rejection occurring under these regimens remains poorly understood.

Alloreactive CD8+ T cells present a major barrier to allograft acceptance as they are major drivers of acute cellular rejection (ACR)^15^. Insights into graft-infiltrating T cells and their clonality during allograft rejection in previous studies were performed with bulk TCR sequencing analyses^16-20^. However, these studies only sequenced TCR? chains, which, in absence of TCRα chain sequencing, do not indicate true clonality nor T cell type (e.g., CD4^+^ or CD8^+^). Similarly, bulk RNA sequencing (bulkseq) has identified rejection-associated transcripts^21-23^, which contributed to the understanding of transplant rejection on the transcriptomic level. However, a major limitation of bulkseq analysis (both TCR and whole transcriptome) is that they do not attribute mRNA transcripts to individual cells, particularly T cells driving allograft rejection.

In contrast to bulkseq, single cell RNA sequencing (scRNAseq) has enabled transcriptomic analysis of individual cells^24^. Additionally, single cell TCR sequencing (scTCRseq) enables assessment of T cell clonality (by single cell pairing of TCRα/β chains) in combination with scRNAseq^25^. Although scRNAseq was used to characterize macrophages in a patient undergoing mixed rejection^26^ and to define donor versus recipient leukocytes in patients undergoing antibody mediated rejection^27^, single cell analysis of transcripts and CD8^+^ T cell clonality in the acutely rejecting renal allograft remains undefined. Another study profiled bronchoalveolar lavage (BAL) fluid cells by scRNAseq from patients undergoing ACR of lung allografts before and after treatment with glucocorticoids^28^. However, analysis of lung tissue during rejection was not performed, so the relationship of BAL fluid-derived T cells to T cells and other immune cells infiltrating lung allograft tissue during rejection remains unclear.

Here, we provide the first combined scRNAseq/scTCRseq analysis of human kidney allograft biopsies from patients undergoing ACR, including analyses of serial biopsies over time and comparative analyses of paired urine and allograft biopsy samples. Our analyses yield important new findings on the nature of human renal allograft rejection including: (i) remarkable restriction of CD8^+^ T cell clonal expansion; (ii) the type of maintenance IS affects gene expression within expanded CD8^+^ T cell clonotypes (CD8_EXP_) observed in index biopsies (biopsies obtained at time of rejection); (iii) persistence of CD8_EXP_ observed in both index and subsequent renal allograft biopsies (even months later), reflecting long-term clonal persistence and adaptation despite rejection treatment; and (iv) correlation of CD8_EXP_ observed in renal allograft rejection biopsies with those obtained in urinary sediment. Together, these results provide novel and fundamental insights into allograft rejection and how CD8_EXP_ respond to anti-rejection therapies. Our results indicate that combined scRNAseq/scTCRseq has the potential to instruct the personalization and enhancement of anti-rejection therapy to improve long-term allograft survival.

## Results

### scRNAseq analysis of acutely rejecting human kidney allografts

To understand ACR at the single cell level, we performed scRNAseq with 5’ V(D)J sequencing on index kidney allograft biopsies obtained from 13 individual participants: 10 biopsies from participants undergoing an ACR episode and 3 control biopsies from participants not experiencing rejection. Hypothesizing that IS type may influence rejection phenotype, ACR samples included biopsies from 4 participants on tacrolimus, 3 on iscalimab, and 3 on belatacept maintenance IS (**Fig. 1A**). Participants varied in terms of age, sex, race, etiology of end-stage renal disease, and donor type (living or deceased) and there were no significant differences in the number of HLA mismatches between IS groups (**Extended Data Fig. 1**). Using our approach to collect and freeze intact biopsies^29,30^ to allow for batch analyses and a novel cold-digestion protocol^31^ to minimize temperature-driven artifacts in gene expression, biopsy-derived cells were subjected to scRNAseq. After alignment, quality control, and integration, uniform manifold approximation and projection (UMAP) analysis of all 13 index biopsies showed that cells were distributed across differing clusters with no clusters completely dominated by any particular sample **(Fig. 1B)**, indicating successful normalization and integration.

**Figure 1.**
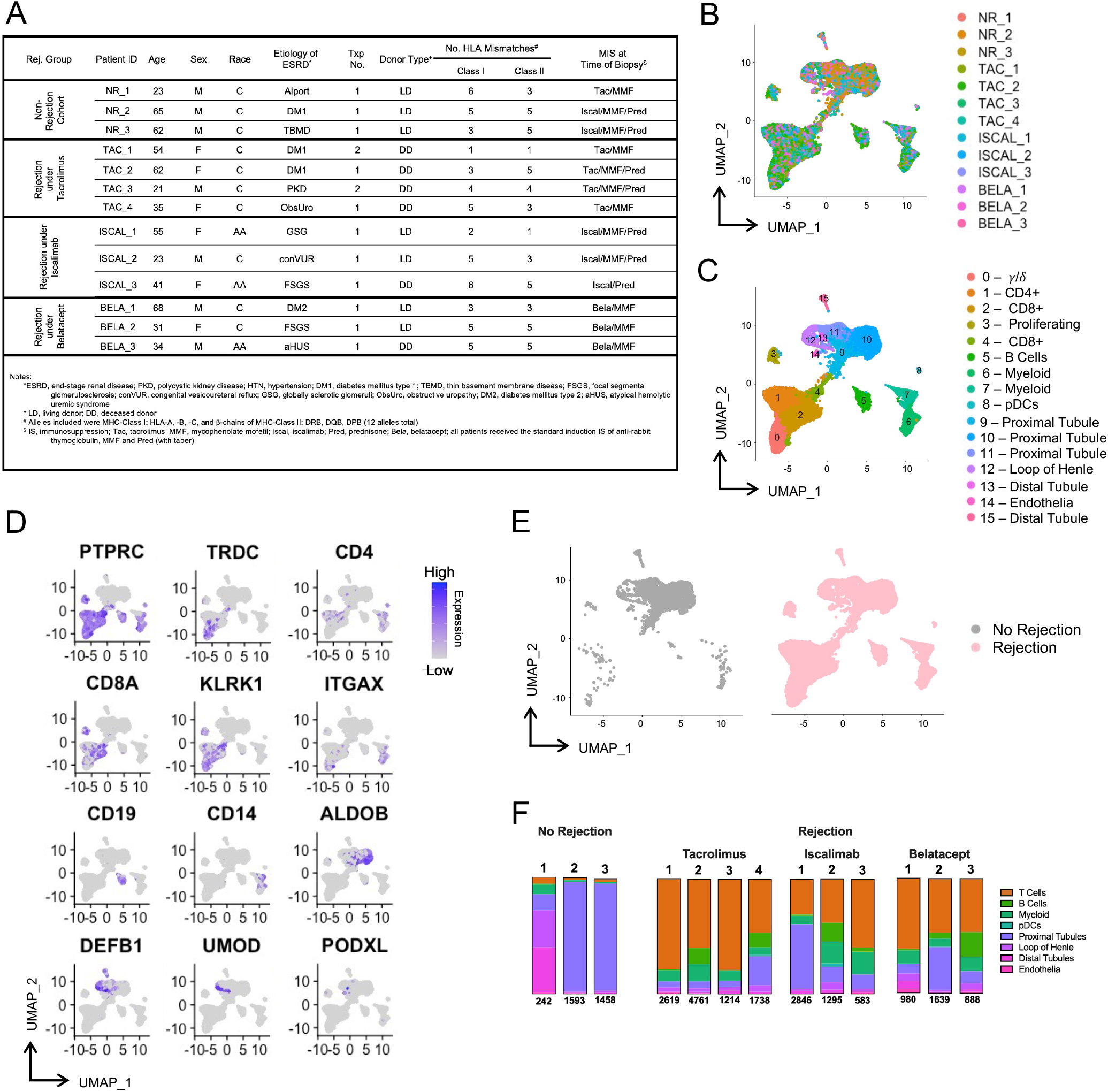
Single cell RNA sequencing analysis of transplanted kidney allografts. Single cell suspensions from thirteen different biopsies (3 without rejection, 10 with rejection) were individually subjected to 5’ single cell RNA sequencing on the 10X platform with V(D)J sequencing. After alignment using Cell Ranger, cells with >25% mitochondrial content and <200 genes, including additional low-quality cells, were removed, and samples were integrated using Seurat. **(A)** Participant demographics of the 13 samples included in index biopsy analyses. **(B, C)** UMAP plots displays cell contribution by sample and cell type (annotations based on differentially expressed genes). **(D)** Expression of “signature” genes across cell types. Blue color intensity reflects the expression level of individual genes within given cells. **(E)** Separation of samples based on rejection status. UMAP plots show cells from non-rejection samples (gray, left plot) versus rejection samples (pink, right plot). **(F)** Frequency of cell types within each sample are displayed in bar graphs.

Initial cluster differentiation revealed 16 clusters of cells, including multiple immune and non-immune, kidney-derived cell populations. Cell types were identified based on differentially expressed genes and expression of canonical markers^32^ (**Supplementary Table 1**). Within the immune cells, we observed ψ/8 (clusters 0), CD4^+^ (cluster 1), CD8^+^ (clusters 2-4), including a population of proliferating CD8^+^ and ψ/8 (cluster 3) T cells; B cells (cluster 5); myeloid cells (clusters 6,7); and plasmacytoid dendritic cells (pDCs) (cluster 8) (**Fig. 1C**). Of the kidney-derived cells, we identified proximal tubule (clusters 9-11), loop of Henle (cluster 12), distal tubule (clusters 13,15), and endothelial (cluster 14) cells (**Fig. 1C**). Individual gene expression plots across clusters were consistent with cell cluster definitions. Immune cell clusters 0-8 expressed *PTPRC*, confirming that all immune clusters were comprised of leukocytes, as well as other cell type-defining markers, including *TRDC* (clusters 0), *CD4* (cluster 1), *CD8A* (clusters 2-4), *KLRK1* (clusters 0-4), *ITGAX* (cluster 0,7,8), *CD19* (cluster 5), and *CD14* (clusters 7,8) (**Fig. 1D**). In humans *CD4* is also expressed by myeloid cells^33^, which we also observed (clusters 7,8) (**Fig. 1D**). As expected, non-rejection biopsies were dominated by kidney-derived cells, while rejection biopsies had prodigious immune infiltrates (**Fig. 1E,F**), consistent with their rejection histology (**Extended Data Fig. 2**).

### CD8^+^ T cells dominate infiltrating immune cell populations in rejecting kidney allografts

To further characterize immune infiltrates, immune cell clusters from the 10 index rejection biopsies were subsetted and re-analyzed. Subsequent cellular annotations revealed: 3 γ/δ T cell clusters, including effector (cluster 0), chronically stimulated (cluster 1), and resident memory (cluster 2) populations; 4 CD8^+^ T cell clusters, including effector (cluster 3), resident memory (cluster 4); memory (cluster 5), and exhausted (cluster 6) populations; 3 CD4^+^ T cell clusters, including follicular helper (cluster 7), memory (cluster 8), and Th17 (cluster 9) cells; 4 myeloid clusters, including macrophages (cluster 11), dendritic cells (cluster 12), extravascular monocytes (cluster 13); and plasmacytoid dendritic cells (pDCs) (cluster 14); 2 B cell populations, including naïve (cluster 15), and class-switched (cluster 16) B cells (**Fig. 2A,B**, **Supplementary Table 2**). Again, both CD8^+^ and γ/δ T cells were present in a proliferating T cell population (cluster 10).

**Figure 2.**
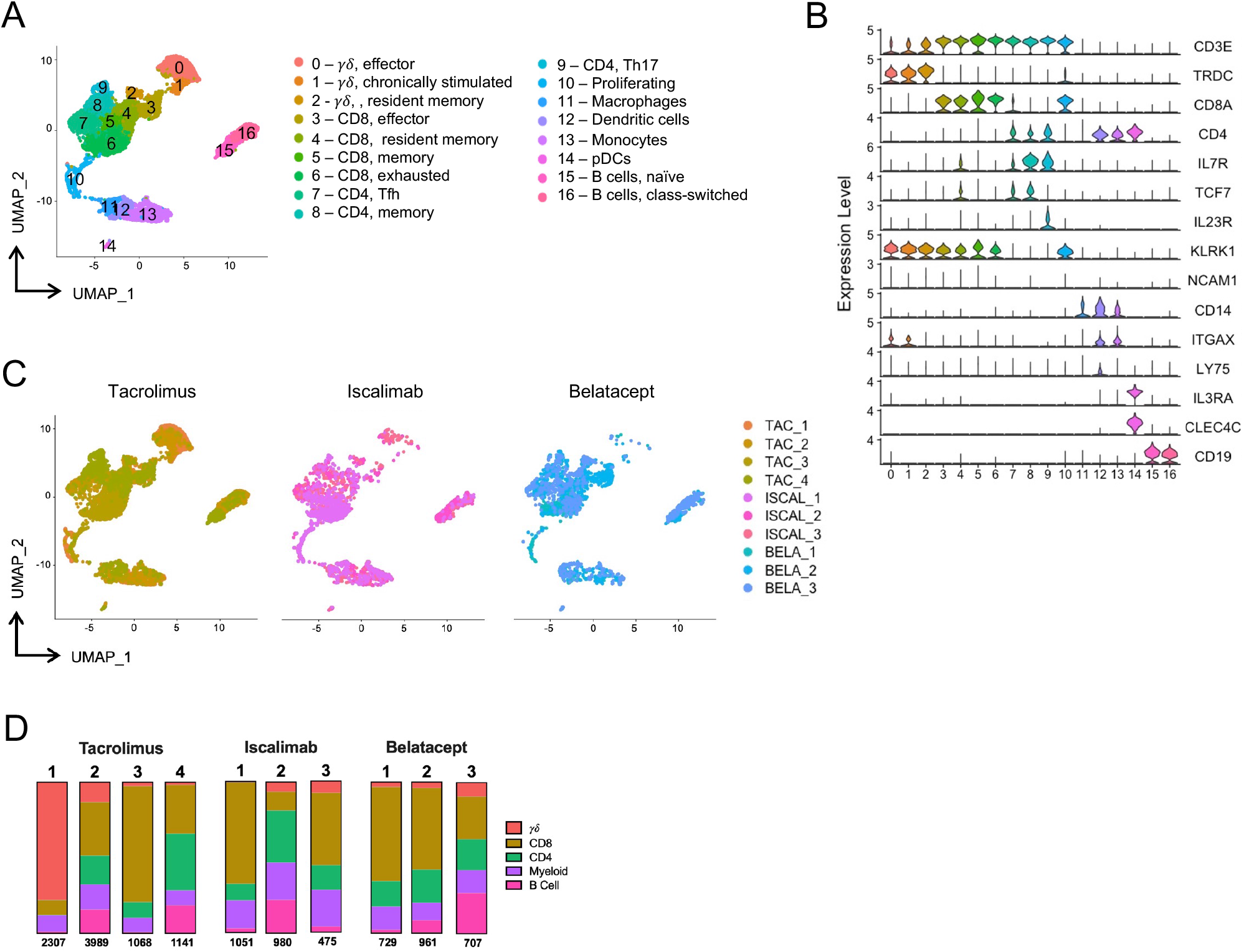
Diverse immune cell infiltrate during kidney allograft rejection. Index samples from the 10 participants undergoing rejection were integrated, clusters annotated as non-immune cells were removed, and the data were re-normalized and re-clustered using Seurat. **(A)** UMAP plot shows immune cell clusters and accompanying annotations. **(B)** Violin plots display the relative gene expression level of indicated genes across each cluster. **(C)** Samples were segregated according to maintenance IS type. UMAP plots shows immune cell clustering of samples from participants with rejection under tacrolimus (left plot, shades of mustard); iscalimab (middle plot, shades of pink); or belatacept (right plot, shades of blue) maintenance immunosuppression **(D)** Frequency of cell types within each sample are displayed in bar graphs.

To examine the influence of maintenance IS regimens on immune cell types present within the allograft, cells were colored according to their IS regimens (tacrolimus [mustard], iscalimab [pink], or belatacept [blue]) **(Fig. 2C**). Although one participant on tacrolimus IS had a dominant influx of γ/δ T cells, most immune cell clusters were present at similar levels for all IS regimens. Overall, maintenance IS type did not grossly affect overall immune cell composition of index biopsies (**Fig. 2D**). Notably, all 10 samples are dominated by T cells, with a majority having more CD8^+^ T cells than CD4^+^ T cells (**Extended Data Fig. 2**).

### CD8_EXP_ gene expression is influenced by IS regimen

As CD8^+^ T cells are primary drivers of ACR, analyses were refocused on just CD8^+^ T cells from index biopsies in the 10 participants with ACR **(Fig. 3A)**. To get greater clarity of the cellular phenotypes associated with each cluster, we compared their differentially expressed genes (**Supplementary Table 3**) as well as expression of markers associated with T cell activation/effector function (*PRF1, GZMB, IFNG, HLA-DRA, CX3CR1, TBX21, MKI67*); exhaustion (*TOX, PDCD1, HAVCR2, LAG3, TIGIT, NR4A1, NKG7*); and resident/circulating memory (*ZNF683, PRDM1, CD69, ITGAE, CXCR6, S1PR1, SELL*) (**Fig. 3B**). For example, in addition to their high-level expression of *PRF1* and *GZMB*, circulating memory cells (CD8_CIRCM_), which includes both effector and central memory cells^35^, in clusters 1 and 2 expressed high levels of *S1PR1* (**Fig. 3A,B**), which promotes their tissue egress^34,35^. Cells in clusters 2 and 4 were defined as resident memory cells (CD8_RM_) based on their expression of *ZNF683, CD69*, and *CXCR6* (**Fig. 3A,B**), which are part of a tissue residency genetic program^36^. In the kidney, not all resident memory cells express *ITGAE*^37,38^. CD8_RM_ were further subdivided based on their differential expression of activation markers *GZMB, IFNG*, and *HLA-DRA* in cluster 4 relative to cluster 2 (**Fig. 3A,B**). Cells in clusters 3, 5, 6, and 7 were likely existing along a continuum of activation (CD8_ACTIV_) and exhaustion (CD8_EXH_) based on their expression of markers associated with exhaustion^39^ (*TOX, HAVCR2, PDCD1, TIGIT, LAG3)* and varying expression of activation/effector function genes (*GZMB* and *IFNG)* (**Fig. 3A,B**). For example, cells in cluster 3, 5, and 7 may be more exhausted as they lack expression of *GZMB* and have lower levels of *IFNG* while cells in cluster 6 are more activated based on their higher expression of *GZMB* and *IFNG* (**Fig. 3A,B**). Finally, cluster 8 represents a population of proliferating CD8^+^ T cells (CD8_PROLIF_) based on expression of *MKI67* and other proliferation-associated genes (**Fig. 3A,B**, **Supplementary Table 3**) Thus, allograft-infiltrating CD8^+^ T cells are heterogeneous with phenotypes consistent with circulating memory, resident memory, and varying states of activation, exhaustion, and proliferation.

**Figure 3.**
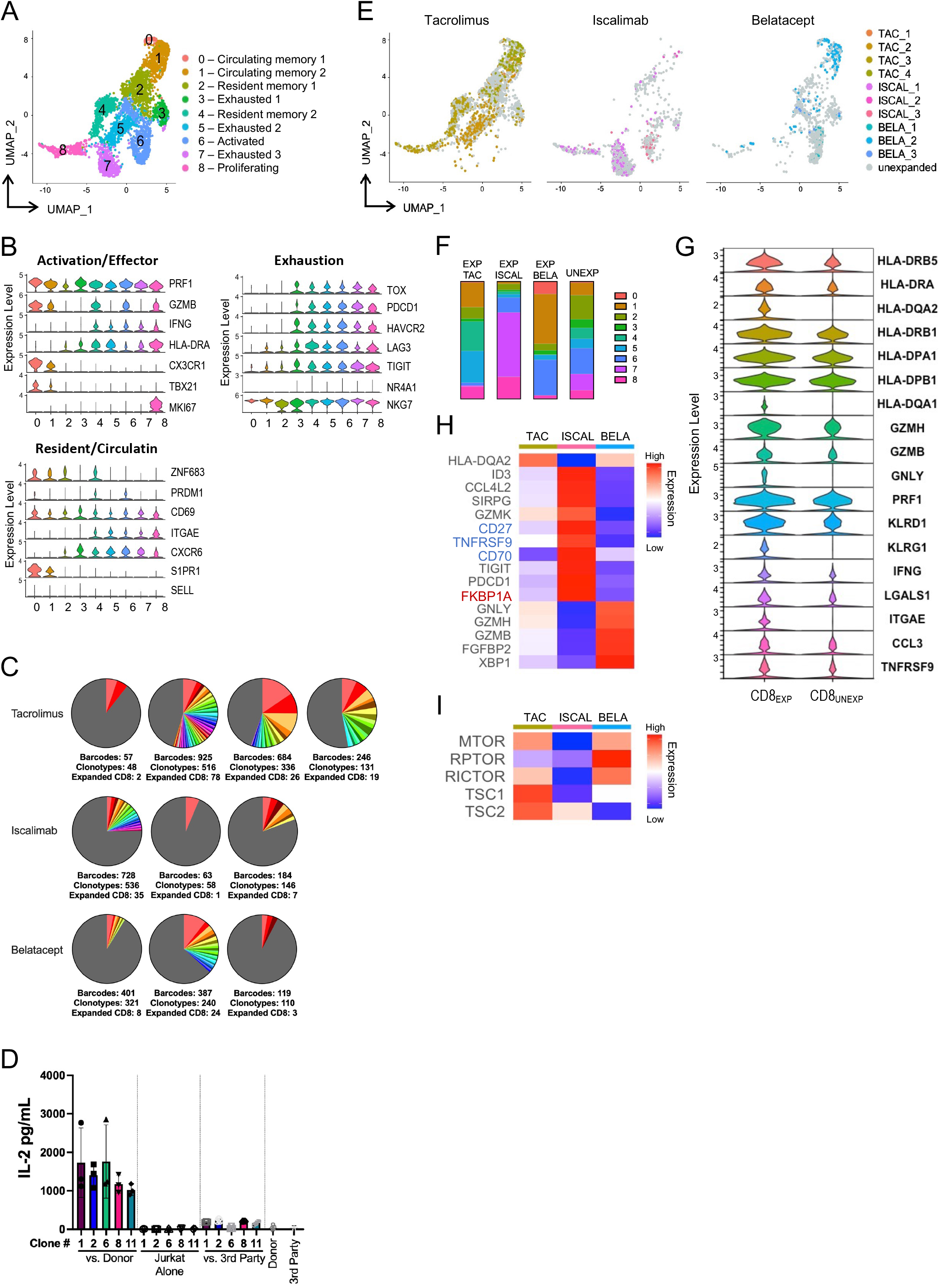
Analysis of infiltrating CD8^+^ T cells in kidney allograft rejection. CD8^+^ clusters from the immune cells analysis were identified for further analyses; CD4^+^ and γ/δ T cells were removed. The samples were then re-normalized and re-clustered using Seurat. **(A)** UMAP plot shows cell type annotations based on differentially expressed genes. **(B)** Violin plots show relative expression level of indicated genes selected to characterize cell cluster phenotypes as activated/effector, exhausted, and resident/circulating memory. **(C)** Pie charts display number and frequency of expanded CD8^+^ clonotypes (CD8EXP) found in the biopsy during rejection by participant sample, based on their unique CDR3α/β sequences. Expanded clonotypes are defined as having >2 cells with identical CDR3 α/β sequences. Different colors represent individual CD8EXP (gray area represents unexpanded clonotypes [CD8_UNEXP_]) and the size of the colored area represents the relative size of the expanded clonotypes. **(D)** Full length TCRs with unique CDR3α/β sequences derived from 5 CD8_EXP_ were subcloned into individual Jurkat76 cells. Individual clones were cultured in triplicate either alone, with donor or with 3rd party T cell-depleted PBMCs for 20 hours and IL-2 levels in the supernatant measured via ELISA. Results show the level of IL-2 in pg/ml for each condition +/-SD done in triplicates. **(E)** Clustering of CD8EXP based on maintenance IS type. UMAP plots show clustering of CD8_EXP_ (colored dots) versus CD8_UNEXP_ (gray dots) from participants under either tacrolimus (left plot, shades of mustard); iscalimab (middle plot, shades of pink); or belatacept (right plot, shades of blue) maintenance IS. **(F)** Bar graphs display the fraction of expanded clonotypes (tacrolimus, iscalimab, or belatacept) and unexpanded clonotypes contributing to each CD8^+^ T cell cluster. **(G)** Violin plots show the relative expression of indicated genes in CD8_EXP_ and CD8_UNEXP_. **(H)** Heatmap displays (average) expression of unsupervised differentially expressed genes in CD8_EXP_ under tacrolimus, iscalimab, and belatacept maintenance IS. Blue text denotes 3 TNF family member genes and red text denotes *FKBP1A*, a target of tacrolimus. **(I)** Heatmap displays a supervised analysis of the average expression of mTOR pathway-related genes in CD8_EXP_ from participants under tacrolimus, iscalimab, and belatacept maintenance IS.

### Limited numbers of clonally expanded CD8^+^ T cells are present in rejecting allografts

CD8^+^ T cell clonality within the rejecting allograft was determined using the 10X Genomics Chromium single cell 5’ V(D)J platform. Full-length CDR3α/β sequences were obtained from ~90% of transcriptionally defined T cells and expanded clonotypes were defined as a CDR3α/β paired sequence present on >2 cells. Strikingly, we found a limited number of expanded CD8^+^ clonotypes (CD8_EXP_) across all 3 IS modalities (average of 20 unique CD8_EXP_ per biopsy), while the majority of CD8^+^ T cells were unexpanded (CD8_UNEXP_) (**Fig. 3C**). Intriguingly, CD4^+^ clonal expansion was minimal, except for participants TAC_4 and ISCAL_2, which had slightly higher proportions of expanded CD4+ T cells (CD4_EXP_) **(Extended Data Fig. 3**). Surprisingly, the level of clonal expansion was not correlated with the number of HLA mismatches, rejection grade, absolute lymphocyte count (ALC), or IS modality **(Extended Data Fig. 4**). While T cells can express 2 TCRα chains, there was no significant difference in the percentage of T cells bearing two TCRα chains between the expanded (7.1%) and unexpanded (8.9%) clonotypes, indicating clonal expansion is driven by antigen recognition by cells expressing a single TCR.

To further understand the donor specificities of the CD8_EXP_, we arbitrarily chose 5 CD8_EXP_ CDR3α/β sequences from the scTCRseq data to subclone into Jurkat76 cells (a thymoma cell line lacking TCRα/β expression)^40^. Resulting Jurkat76 transfectants were cultured with T cell-depleted PBMCs from the recipient’s kidney donor or 3^rd^ party cells and assayed for responses via IL-2 production. Strikingly, all 5 clones responded to donor, but not 3^rd^ party, cells (**Fig. 3D**), confirming the alloreactivity of those CD8_EXP_ identified in the rejecting kidney biopsy.

### Maintenance IS type affects CD8_EXP_ gene expression

Interestingly, cluster distribution of CD8_EXP_ varied based on maintenance IS. CD8_EXP_ from tacrolimus-treated participants were distributed across all clusters but were more frequently represented in the CD8_RM_ and CD8_EXH_ populations (clusters 4, 5) than CD8_EXP_ from belatacept-or iscalimab-treated participants (**Fig. 3E,F**). CD8_EXP_ from iscalimab-treated participants predominantly resided in another CD8_EXH_ population (cluster 7) and the CD8_PROLIF_ population (cluster 8), which was distinct from CD8_EXP_ from belatacept-or tacrolimus treated participants (**Fig. 3E,F**).. CD8_EXP_ from belatacept-treated participants clustered predominantly in both the CD8_CIRCM_ (cluster 1) and CD8_ACTIV_ (cluster 6) populations. CD8_UNEXP_ cells were most represented in clusters with the lowest levels of CD8^+^ T cell activation (CD8_CIRCM_, CD8_RM_, and CD8_EXH_) (**Fig. 3E,F**). Further examination of the differential gene expression between CD8_EXP_ and CD8_UNEXP_ revealed that CD8_EXP_ had higher expression of *HLA* markers, activation and effector markers (*GZMH, GZMB, GNLY, PRF1, KLRD1, KLRG1, IFNG, ITGAE*), chemokines (*CCL3, CCL4*) and chemokine receptors (*CCL4L2*), and TNF family members *(TNFRSF9*) **(Fig. 3G**). Thus, CD8_EXP_ express alloreactive TCRs and have gene expression consistent with cells that have undergone TCR-mediated activation.

We next examined the differentially expressed genes (DEGs) in CD8_EXP_ between various IS. Iscalimab CD8_EXP_ had increased expression of TNF family members such as *CD27, TNFRSF9*, and *CD70* (blue), as well as *FKBP1A* (red), an intracellular tacrolimus binding protein. Interestingly, *FKBP1A* expression is decreased in CD8_EXP_ under tacrolimus and belatacept IS. **(Fig 3H**). In contrast, belatacept CD8_EXP_ show upregulation of activation markers such as *GNLY, GZMH*, and *GZMB* when compared to iscalimab and tacrolimus CD8_EXP_.

Previously, our group demonstrated increased mTOR activity in peripheral blood CD8^+^ T cells in patients with ACR under belatacept, but not tacrolimus, and treatment of belatacept-refractory ACR with everolimus mitigated their ACR^5^. Based on this, we performed a supervised analysis of mTOR pathway-related genes in the CD8_EXP_ under the 3 maintenance IS regimens (**Fig 3I**). Notably, CD8_EXP_ from participants under belatacept IS showed not only a significantly increased expression of mTOR complex genes (*mTOR, RPTOR, and RICTOR*), but also decreased expression of 2 negative regulators of mTOR activation (*TSC1, TSC2*). Combined, these data indicate that rejections arising under differing IS are associated with varying gene expression of potential therapeutic targets.

### CD8_EXP_ clonal populations may expand, contract, or persist in response to anti-rejection treatment

We next examined the persistence of CD8_EXP_ clonal populations after anti-rejection therapy. Participant TAC_3 experienced an index Banff ACR 1B rejection on post-transplant day (PTD) 217, which was treated with rabbit anti-thymocyte globulin (rATG) and prednisolone (**Extended Data Figure 5**). Two weeks later (PTD 232), a follow-up biopsy revealed histologic improvement to Banff borderline lesion and a substantial decrease in the total numbers of CD8^+^ T cells. However, scRNAseq analysis revealed no significant change in the frequency of CD8_EXP_ from 10.7% (26 out of 336 total clonotypes) in the index biopsy to 13% (24 out of 182) (**Fig. 4A**). Interestingly, out of the 24 CD8_EXP_ that were identified at PTD 232, 10 were identical to those present as CD8_EXP_ in the index biopsy (PTD 217) (**Fig. 4A**). Rejection treatment resulted in marked reduction in CD8_EXH_ (cluster 4), CD8_ACTIV_ (cluster 5) and CD8_PROLIF_ (cluster 6) CD8_EXP_ (**Fig. 4B-D, Supplementary Table 4**). Notably, at PTD 232, a dominant CD8_EXP_ population with a CD8_RM_ phenotype (cluster 0) appeared. Nine weeks later (PTD 295), another biopsy revealed no histologic rejection, yet scRNAseq analysis revealed 5 distinct CD8_EXP_, 3 of which were from previous biopsies (**Fig. 4A,D**). DEG analysis between the CD8_EXP_ from the 3 timepoints revealed that clonotypes from the index rejection biopsy expressed effector function genes (*GZMB, GZMK, GNLY)* but also exhaustion markers (*LAG3, HAVCR2, TIGIT)*, while clonotypes two weeks later (PTD 232) displayed resident memory genes (*ZNF683, CD69, ITGAE)* and clonotypes two months later (PTD 295) expressed *GZMK, LRB1*, as well as chemokines *XCL1, XCL2* (**Fig. 4E**). Thus, even though histologic rejection resolved, CD8_EXP_ persisted at 11 weeks after initial rejection despite rATG and corticosteroid anti-rejection treatment.

**Figure 4.**
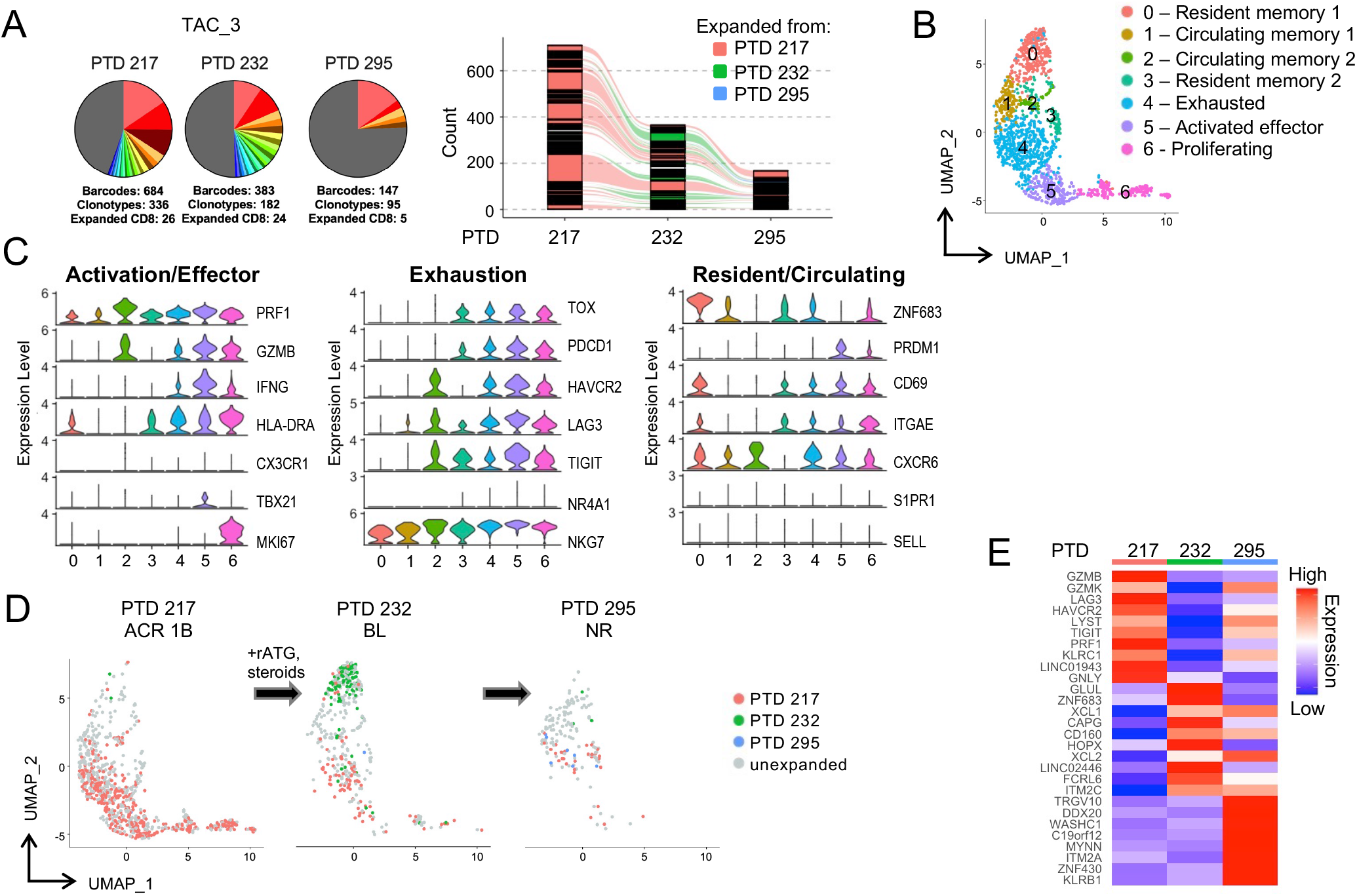
Temporal scRNAseq analysis of the response to anti-rejection therapy under tacrolimus maintenance immunosuppression (IS). A participant on tacrolimus IS (TAC_1) was diagnosed with ACR 1B on PTD 217 and a biopsy was obtained prior to anti-rejection treatment with rATG and CCS. A second biopsy was obtained on PTD 232 and the participant was diagnosed with borderline rejection. A third biopsy was taken at PTD 295 and the participant was diagnosed with no rejection. **(A)** Pie charts display number and frequency of expanded clonotypes found in the index biopsy (PTD 217) and subsequent follow-up biopsies (PTD 232, PTD 295). Bar graph shows overlapping clonotypes across the 3 timepoints. **(B)** UMAP shows CD8^+^ clusters in an integrated analysis of all timepoints. **(C)** Violin plots show relative expression level of indicated genes selected to characterize cell cluster phenotypes as activated/effector, exhausted, and resident/circulating memory. **(D)** Temporal analysis of CD8_EXP_ following anti-rejection therapy. UMAP plots show clustering of expanded (CD8_EXP_, colored dots) versus unexpanded (CD8_UNEXP_, gray dots) CD8^+^ clonotypes from the participant at PTD 217 (left plot); PTD 232 (middle plot); or PTD 295 (right plot). CD8_EXP_ first expanded on PTD 217 are shown in pink, those first expanding PTD 232 are shown in green, and those first expanding PTD 295 are shown in blue. **(E)** Heatmap shows average expression of unsupervised differentially expressed genes found between CD8_EXP_ at each timepoint.

Participant ISCAL_1 presented with a Banff ACR 2A rejection under iscalimab-based IS at one-month post-transplant that was treated with rATG and a corticosteroid taper for persisting Banff 1B rejection. A biopsy obtained on PTD 60 revealed a Banff 1A rejection (**Extended Data Figure 5**), and scRNAseq analysis revealed 35 individual CD8_EXP_, most of which resided in clusters 0 (resident memory), 4 (exhausted), and 5 (proliferating) (**Fig. 5B,C**, **Supplementary Table 5**). Tacrolimus-based anti-rejection treatment was initiated, and a biopsy obtained 2.5 weeks later (PTD 78) demonstrated no rejection and significantly reduced CD8_EXP_ in clusters 4 (exhausted) and 5 (proliferating). However, there was no change in the frequency of CD8_EXP_ (6.5% at the first timepoint and 6.3% at follow-up), and nearly half of the CD8_EXP_ were identical to those in the first biopsy (**Fig. 5A,D**). Strikingly, at this time, a majority of CD8_EXP_ resided in a new exhausted population (cluster 3) (**Fig. 5B,C**, **Supplementary Table 5**). Following rejection resolution, tacrolimus was tapered. A repeat biopsy obtained approximately 1 year later (PTD 336) revealed histologic resolution of rejection, and only 1 CD8_EXP_ persisted from the index biopsy (**Fig. 5A,D)** which had a resident memory phenotype (evidenced by *ZNF683* expression) (**Fig. 5E)**. Notably, the TCR expressed by this clone was one of the TCRs defined as alloreactive **(Fig. 3D)**. This demonstrates that, despite resolution of rejection, alloreactive CD8^+^ T cell clones can persist for at least a year.

**Figure 5.**
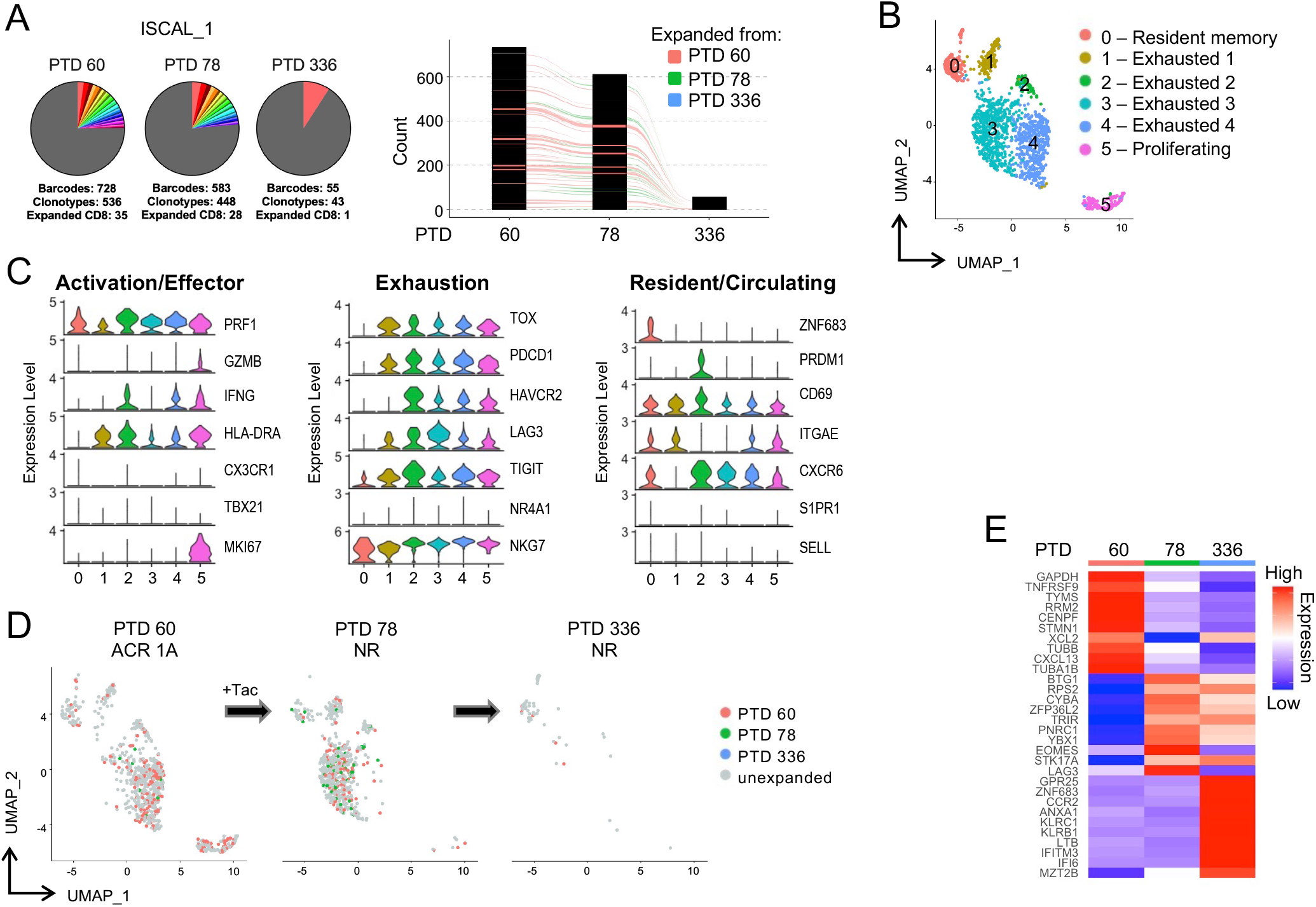
Temporal scRNAseq analysis of the response to anti-rejection therapy under iscalimab maintenance immunosuppression (IS). A participant on iscalimab IS (ISCAL_1) was diagnosed with ACR 1A on PTD 60 and a biopsy was obtained prior to anti-rejection treatment with tacrolimus. A second biopsy was obtained on PTD 78 and the participant was diagnosed with no rejection. A third biopsy was taken at PTD 336 and the participant was again diagnosed with no rejection. **(A)** Pie charts display number and frequency of expanded clonotypes found in the index biopsy (PTD 60) and subsequent follow-up biopsies (PTD 78, PTD 336). Bar graph shows overlapping clonotypes across the 3 timepoints. **(B)** UMAP shows CD8^+^ clusters in an integrated analysis of all timepoints. **(C)** Violin plots show relative expression level of indicated genes selected to characterize cell cluster phenotypes as activated/effector, exhausted, and resident/circulating memory. **(D)** Temporal analysis of CD8_EXP_ following anti-rejection therapy. UMAP plots show clustering of expanded (CD8_EXP_, colored dots) versus unexpanded (CD8_UNEXP_, gray dots) CD8^+^ clonotypes from the participant at PTD 60 (left plot); PTD 78 (middle plot); or PTD 336 (right plot). CD8_EXP_ emerging on PTD 60 are shown in pink, on PTD 78 are shown in green, and those emerging on PTD 336 are shown in blue. **(E)** Heatmap shows average expression of unsupervised differentially expressed genes found between CD8_EXP_ at each timepoint.

A third participant, treated with iscalimab-based maintenance IS (ISCAL_3), was diagnosed with a Banff 1B rejection at 20 weeks post-transplant (PTD 137) (**Extended Data Figure 5**). scRNAseq analysis revealed 7 distinct CD8_EXP_, most of which had gene expression consistent with an activated effector phenotype (**Fig. 6A,D-F**, **Supplementary Table 6**). Interestingly, scRNAseq analysis of urine sediment at the same timepoint contained 11 CD8_EXP_, 4 of which were identical to the CD8_EXP_ in the biopsy (**Fig. 6B,C**). To treat their rejection, the participant was converted to tacrolimus IS, given a prednisolone pulse, and iscalimab was discontinued. Two weeks later (PTD 151), a repeat biopsy revealed Banff ACR 1B rejection and an increase in the CD8_EXP_ clonal frequency (4.7% to 7.2%). Importantly, each CD8_EXP_ present in the index biopsy were also observed in the second biopsy, and 16 CD8_EXP_ were found in both the biopsy and urine samples **(Fig. 6A,C)**. Most CD8_EXP_ from this biopsy were identified as CD8_CIRCM_ and CD8_ACTIV/EFF_ cells (clusters 1,3-5) (**Fig. 6D-F**). Mycophenolate mofetil (MMF) was added to the maintenance IS regimen and four weeks later (PTD 179), repeat biopsy revealed a borderline lesion, with a continued persistence of CD8_EXP_ (6.3%) **(Fig. 6A)**. The majority of CD8_EXP_ at this timepoint were identified as CD8_EXH_ cells (cluster 2) (**Fig. 6D-F**). Approximately 72% (13 out of 18) CD8_EXP_ in the PTD 179 biopsy had been observed in prior biopsies and a similar persistence of CD8_EXP_ was observed in the urine (**Fig. 6A,B**). An approximate 50% overlap in CD8_EXP_ were noted in both biopsy and the urine at PTD 179 (**Fig. 6C**). Four months later (PTD 291), the participant was diagnosed with a Banff 1B mixed acute rejection. scRNAseq again demonstrated persistence of CD8_EXP_ at a frequency 9.5% of all clonotypes (16 out of 168) and roughly half of these were observed in earlier biopsies. Interestingly, also at this timepoint, a new clonotype appeared and localized to cluster 0, a cluster not present in the earlier samples and whose gene expression profile was consistent with a pathogenic resident memory phenotype (**Fig. 6F,G**). Taken together, these data show that, during unresolved rejection, treatment with tacrolimus, corticosteroids, and MMF failed to eliminate CD8_EXP_ instead drove substantial changes in their gene expression.

**Figure 6.**
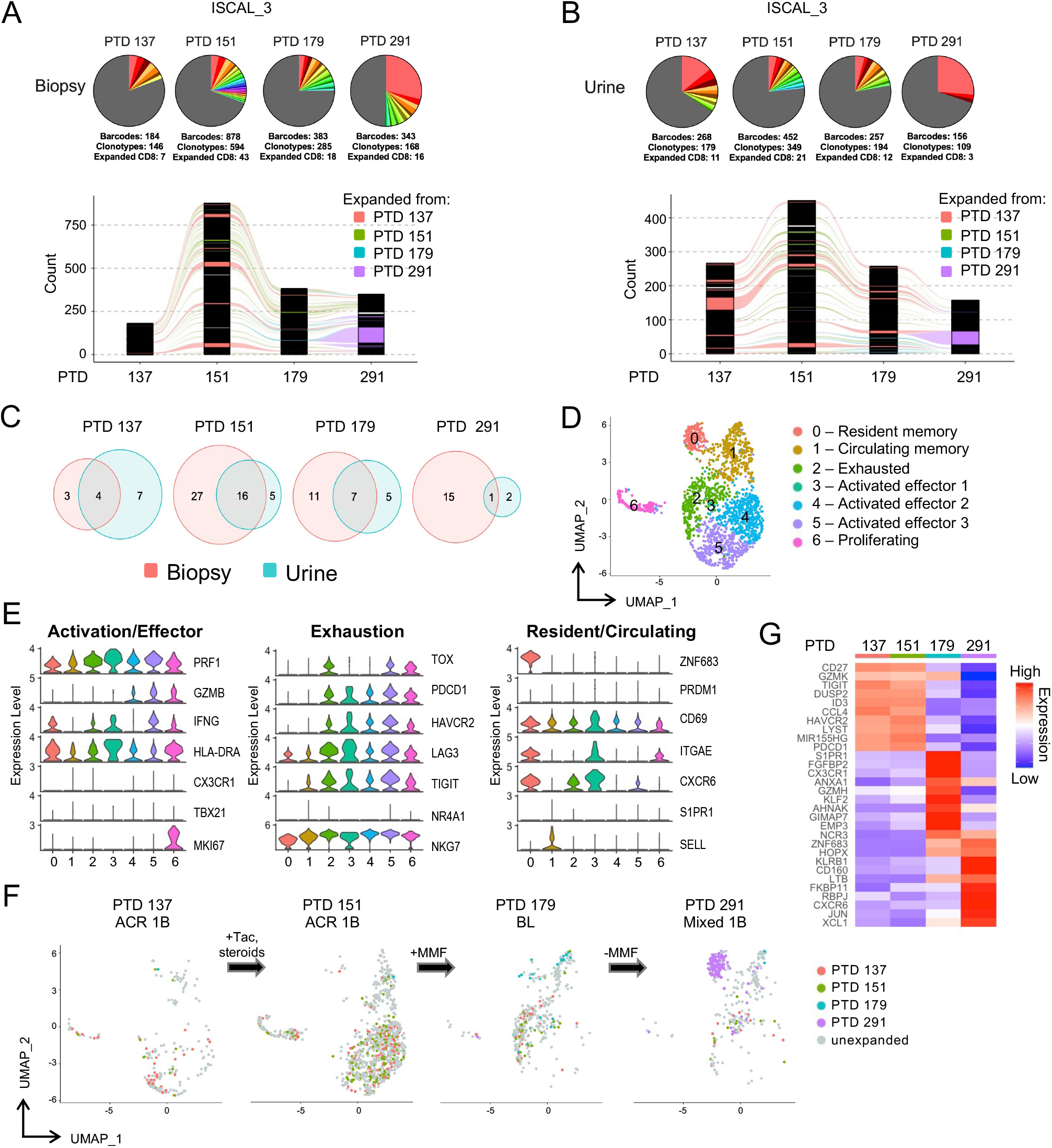
Temporal scRNAseq analysis of CD8EXP in biopsy and paired urine samples in a participant undergoing treatment-refractory rejection. A participant on iscalimab immunosuppression (ISCAL_3) was diagnosed with ACR 1B on PTD 137 and a biopsy was obtained prior to anti-rejection treatment with tacrolimus conversion and CCS. A second biopsy was obtained on PTD 151 and the participant was diagnosed with ACR 1B. MMF was then added to the anti-rejection regimen. A third biopsy was taken at PTD 179 and the participant was diagnosed as borderline and MMF was tapered off. A final biopsy was taken at PTD 291 and showed mixed 1B rejection. **(A, B)** Pie charts (top) display number and frequency of expanded clonotypes found at each biopsy **(A)** and urine **(B)** sample and bar graphs (bottom) display clonotypes found at the indicated timepoint. Different colors represent individual expanded clonotypes (gray area represents unexpanded clonotypes) and the size of the colored area represents the relative size of the expanded clonotypes. **(C)** Venn diagrams display overlap of individual CD8_EXP_ clonotypes between biopsies and their paired urine sample at the indicated timepoints. **(D)** CD8^+^ T cells from all timepoints were integrated and renormalized and UMAP plot shows individual CD8^+^ clusters based on differentially expressed genes. Note that some clusters are unique to individual timepoints. **(E)** Violin plots show relative expression level of indicated genes selected to characterize cell cluster phenotypes as activated/effector, exhausted, and resident/circulating memory. **(F)** Temporal analysis of CD8EXP during treatment-refractory rejection therapy. UMAP plots show clustering of expanded (CD8_EXP_, colored dots) versus unexpanded (CD8_UNEXP_, gray dots) CD8^+^ clonotypes from the participant at PTD 137 (left plot); PTD 151 (middle left plot); PTD 179 (middle right plot); or PTD 291 (right plot). CD8_EXP_ emerging on PTD 137 are shown in pink, on PTD 151 are shown in green, PTD 179 are shown in blue, and those emerging on PTD 291 are shown in purple. **(G)** Heatmap shows average expression of unsupervised differentially expressed genes found between expanded clonotypes at each timepoint.

## DISCUSSION

Using single cell genomics, we uncovered an unexpectedly small number of CD8_EXP_ present in renal allograft biopsies undergoing rejection. This seemingly contrasts prior data using bulk TCRϕ3 sequencing approaches of kidney allograft biopsies^16,17,20,41^. We envision a few explanations for these differences. First, transplant patients are often lymphopenic due to induction therapy, which could severely limit the available repertoire. Although we found that patients with a normal absolute lymphocyte count had low numbers of CD8_EXP_, it remains possible that induction therapy drives a long-lasting reduction of the available repertoire. Second, while it is possible that many CD8_UNEXP_ are alloreactive, we think this unlikely as our longitudinal analysis showed that very few CD8_UNEXP_ ended up as CD8_EXP_ in subsequent biopsies. Third, alloresonses may involve immunodominance mechanisms. For example, perhaps the nature of the T cell (i.e., pre-existing memory or TCR avidity) allows a limited number of clonotypes to outcompete for resources and dominate the response. Further work determining TCR specificity of CD8_EXP_ and their presence in naïve and memory T cell populations prior to transplantation will help distinguish between these possibilities.

In this regard, our platform of subcloning of TCRs into Jurkat76 cells and testing them against donor versus third-party T cell-depleted PBMCs will be useful to study the biology of allospecific TCRs. In addition to linking TCR specificity to cell phenotype, screening combinatorial peptide libraries (CPLs) will enable the identification of peptides bound to donor HLA recognized by allospecific TCRs. This is important because a lack of an ability to track and monitor allospecific T cells has been a major impediment to understanding their development and function. Further, as many have suggested, a large fraction of allospecific T cells may be pre-existing memory cells with specificities to pathogens previously encountered^42,43^. Similarly, use of recipient PBMCs and CPLs will allow for identification of TCRs with potential cross-reactivity. As such, future extensions of these studies have the potential to yield insights into the fundamental nature of allorecognition, whether alloreactive TCRs focus on HLA epitopes versus peptide, and protein structure/function analysis of TCR CDR3/HLA/peptide interactions.

Our data also showed that the type of IS affected the gene expression and phenotype of CD8_EXP_. Importantly, scRNAseq provided the resolution to identify potential therapeutic targets that may be exploited for optimal anti-rejection therapy. Our finding that expression of components of the mTOR pathway were elevated in BRR is consistent with our prior data successfully treating patients with BRR^44^. Likewise, increased expression of *FKBP1A* in iscalimab-refractory rejection suggests that such rejections may be sensitive to tacrolimus. Thus, such analysis could provide physicians with informed and personalized approaches to optimize anti-rejection therapies.

We also found that CD8_EXP_ can persist in the kidney allograft for months, despite successful anti-rejection therapy, confirming and extending prior MLR studies in peripheral blood^16^. Interestingly, we also found that individual CD8_EXP_ surviving rejection therapy adapt, and possibly survive, by altering gene expression (i.e., adopting a CD8_RM_ phenotype) in response to anti-rejection therapy. This incomplete CD8_EXP_ elimination and persistence may underlie recurrent rejection and/or long-term smoldering allograft injury, contributing to rejection-associated reduction in allograft survival. Additional work is required to determine the persistence and specificity of TCRs of CD8_EXP_ from participants undergoing anti-rejection therapy and whether these CD8_RM_ cells are associated with protective^45^ versus pathologic^46^ responses. An intriguing point remains – whether complete elimination of CD8_EXP_ will significantly improve the poor allograft survival rates that are observed following treatment of moderate and severe ACR. Addressing this issue may represent a major advance in rejection therapy.

Our observation that clonally identical CD8_EXP_ are found in the urine and biopsy is likely a result of CD8^+^ T cell killing of renal tubular epithelial cells, underlying the predominant histologic feature of ACR (tubulitis). In tubulitis lesions, CD8_EXP_ likely traverse the renal tubular basement membrane and gain access to the urinary space, in contrast to those remaining in the interstitial space. We are currently determining the number and gene expression of CD8_EXP_ in the urine and whether this correlates with the degree of tubulitis. Interestingly, for participant ISCAL_3, the CD8_EXP_ with a CD8_RM_ phenotype was present only in the biopsy (not urine), suggesting their retention in the kidney precluded their ability to traverse the basement membrane.

While we focused on CD8^+^ T cells, many other cell types are amenable to similar analysis, including innate cell populations and kidney-derived cell populations, and both approaches are currently underway utilizing the same samples described herein. In addition, other single cell genomics-based approaches, including spatial transcriptomics (ST) and ATACseq. ST will place the cell populations identified by scRNAseq in their histologic context, while ATACseq may further refine and expand our knowledge of gene regulatory networks active in various cell types.

In summary, scRNAseq analysis of human renal allograft rejection reveals highly restricted CD8_EXP_ that exhibit alloreactivity and varying responses to rejection therapy, including persistence despite rejection resolution. Importantly, these CD8_EXP_ vary in gene expression based on the nature of maintenance IS. These novel and fundamental insights delineate approaches for developing innovative individualized rejection therapies that can be tested in carefully designed clinical trials.

## METHODS

### Participant Enrollment

All participants were independently enrolled in this mechanistic rejection study approved by the University of Cincinnati Institutional Review Board (IRB 2017-4696, 2019-0469). One group of patients were simultaneously enrolled in the CIRRUS I trial (NCT03663335). Enrollment occurred at two Cincinnati, Ohio hospitals (University of Cincinnati Medical Center or The Christ Hospital) upon scheduling of a for-cause renal allograft biopsy. All participants provided informed consent before study procedures occurred, with continuous consent ensured throughout participation.

Tissue samples were collected directly during the biopsy procedure using an 18g biopsy needle and placed immediately in HypoThermosol FRS Preservation Solution (HTS; BioLife Solutions Inc, 101102) on ice. When applicable, clean-catch urine samples were collected and spun down at 300xg to obtain the cellular components. Both sample types were frozen at -80°C in CryoStor CS10 (BioLife Solutions Inc, 210102) in a Mr. Frosty Freezing Container (Nalgene 5100-0001), then stored in liquid nitrogen until ready for analysis.

### Tissue Dissociation

Tissue dissociation protocol was modified from a previously described cold-active protease digestion^31^. Kidney core biopsies were slowly thawed, cut into 1-2mm pieces, then subject to gentle cold digestion on ice using 10% FBS-supplemented RPMI media containing 100mg/mL trypsin inhibitor from soybean (Roche, 10109886001), 10mg/mL collagenase A from clostridium histolyticum (Roche, 10103586001), 10mg/mL collagenase type IV from clostridium histolyticum (Worthington, LS004186), 250U DNase I (Roche, 4536282001), and 5mM CaCl_2_. Digestion was performed twice at 10 minutes each, with intermittent rotating and gentle pipet mixing with wide orifice tips. The digested tissue was passed through a pre-primed 30mm cell strainer, breaking up any remaining visible tissue using a rubber syringe plunger. The single cell suspension was passed through a second pre-primed 30mm cell strainer, then centrifuged at 300xg at 4°C for 5 minutes. Viability and live cells were counted by trypan blue exclusion then resuspended at 1000 cells/mL, per the 10X Genomics Chromium protocol. The cells, kept on ice, were immediately prepared for single-cell barcoding.

### Single cell barcoding, cDNA synthesis and library preparation

All samples were processed for single-cell sequencing following the Chromium Next GEM Single Cell V(D)J Reagent Kits v1.1 protocol. Briefly, cells were uniquely barcoded by using 10X fluidics (10X Genomics Chromium Single Cell Controller) to combine each individual cells with an individual barcoded Single Cell 5’ Gel Bead creating a Gel Beads-in-emulsion (GEMs) solution (10X Genomics, PN-1000165 and PN-1000120). An average of 17,400 cells were loaded to achieve an estimated 10,000 cell recovery. GEM gel beads were dissolved, and cDNA was synthesized from the resulting tagged mRNA transcripts over 14 amplification cycles. 50ng of cDNA was used for the construction of each library. Total gene expression libraries (PN-1000020) and libraries of enriched TCR sequences (PN-1000005) were created using the Single Index Kit Set A (PN-1000213).

### Sequencing, alignment, and generation of matrices

Total gene expression and TCR sequence-enriched libraries were sequenced on the NovaSeq 6000 sequencer using S1, S2 or S4 flow cells, with the goal to obtain >320M reads per sample. Raw base call files were de-multiplexed with Cell Ranger (version 6.1.2) using mkfastq [1]. Reads were aligned to human reference genome (version GRCh38) and gene expression quantified against GENCODE (release 32) using the count function of CellRanger. Genomics data will be made available via the GEO Database, accession number in process.

### Single-cell RNASeq analysis pipeline

Single-cell analysis was carried out with R (version 4.2.0) running inside RStudio (version 4.1.1) using Seurat (version 4.1.0)^47,48^. Cells expressing more than 25% mitochondrial gene transcripts or fewer than 200 genes, including additional low-quality cells, were excluded from the analysis. TCR alpha and beta gene variants were collapsed as singular ‘tcr-alpha’ and ‘tcr-beta’ genes, respectively. Gene expression counts were normalized with the NormalizeData function in Seurat. The samples were integrated using FindIntegrationAnchors and IntegrateData functions from Seurat. This integrated data set was used for principal component analysis, variable gene identification, Shared Nearest Neighbor (SNN) clustering analysis, and creation of UMAP. Metadata was updated to include identities of TCR clonotypes and those categorized as expanded and unexpanded, as described below. Differentially expressed genes were determined using FindMarkers, with a logfc threshold of 1 and minimum percent expression of 0.2. Genes that were differentially expressed at an adjusted p-value < 0.05 were used for analyses.

### TCR clonal analysis

Cell Ranger outputs for the TCRα/µ sequencing data were merged into the Seurat metadata for various integrated analyses. Filtered_contig_annotations.csv and clonotypes.csv files were used to obtain CDR3α/µ information linked to individual barcodes, which were then merged with barcodes from the Seurat metadata to combine scRNAseq analysis with scTCRseq analysis. CD8^+^ cells were identified from the immune cell populations through subsetting of CD8^+^ clusters and removal of cells expressing *CD4, TRDC*, and *CD68*. Clonotypes with identical CDR3α/µ sequences present in >2 cells (identified through unique barcodes) were determined to be expanded. Clonotypes with 2 CDR3? chains or only an individual CDR3α or CDR3? chain were classified as unexpanded. Analysis of the merged Seurat metadata allowed determination of numbers of CD8+ barcodes, total numbers of clonotypes, and numbers of CD8_EXP_ and CD8_UNEXP_ clonotypes.

To determine the position of CD8_EXP_ clonotypes on UMAPs, expanded clonotypes were recalled using their clonotype_id and set as a new identity on the plots. Overlapping clonotypes between biopsy and urine samples were done using the package VennDiagram, and clonotype tracking over sequential timepoints was done using the package Immunarch^49^ v0.9.0 with modified input files to reflect only CD8^+^ cells in the analysis.

### TCR-expressing Jurkats

TCRα/β^-/-^ Jurkat 76 cell lines were provided by Dr. Michael Nishimura (Loyola University). Jurkat76 cells stably expressing the TCRs of interest were generated by transfection using the Neon Transfection System (Thermo Fisher). The cells were transfected with the pCMV(CAT)T7-SB100 plasmid and the pSBbi-Neo Sleeping Beauty vector containing the full length TCR *α* and *β* chains separated by the P2A self-cleaving peptide as previously described^50^. After electroporation, the cells were maintained in RPMI media (RPMI 1640 with 10% FBS, 100 units/mL penicillin, and 100 μg/mL streptomycin). and were incubated at 37°C and 5% CO_2_. Cells expressing the TCR of interest were selected for by using media containing 1.2mg/mL of G418 (Geneticin™, ThermoFisher 10131027). Cells were analyzed and sorted for transfection efficiency via flow cytometry analysis using anti-human CD3 APC/Cy7-conjugated antibody and anti-human CD34 PE-conjugated antibody for TCR-expressing cells (BioLegend, 300426 at 10 μg/mL and BioLegend, 343606 at 1.25 µg/mL).

### Cell Culture and ELISAs

TCR-expressing Jurkat76 cells were cultured overnight (20 hours) in PMA-supplemented non-selection RPMI media at a 1:1 ratio with either T cell-depleted PBMCs derived from the participant’s donor or derived from a third-party healthy donor. Co-culture supernatant was collected following completion of culture and analyzed for IL-2 using the ELISA MAX™ Deluxe Set for human IL-2 (Biolegend, 431804) at a dilution of 1:5.

### Statistics

Statistical analyses, including one-way ANOVA analyses, were done using GraphPad Prism version 9.3.1 with a significance at p<0.05.

## Acknowledgements

We graciously thank all the participants, participant families, and other participant support persons that have made these studies possible. We would like to acknowledge the University of Cincinnati (UC) Transplant Research Team and the UC and Christ Hospital Transplant Care Team for their hard work in consenting, following, and caring for research participants. We acknowledge the UC Pathology team, in particular Dr. Paul Lee, for their contributions to the pathological and histological analyses. We thank Dr. Steve Potter and Andrew Potter for their help with development of the cold protease digestion protocol. We would also like to thank the UC/CCHMC Center for Transplant Immunology for their guidance and support in this project.

This research was also made possible, in part, using the Cincinnati Children’s Single Cell Genomics Core [RRID:SCR_022653], DNA Sequencing and Genotyping Core [RRID:SCR_022630], and Biomedical Informatics Core. We specifically acknowledge the assistance of Kelly Rangel and Shawn Smith from the Single Cell Genomics Core. This work was supported by a grant from Novartis and from Public Health Service grants AI167482 (T.S.), AI142264 (D.A.H, E.S.W.), UH2AR067688 (D.A.H., E.S.W.), UL1TR000077 (D.A.H., E.S.W.), AI169863 (D.A.H., E.S.W., and B.M.B.)

## References

1. D. W, Wiseman. Long-Term Immunosuppression Management: Opportunities and Uncertainties. Clinical journal of the American Society of Nephrology : CJASN. 2021 Aug 2021;16(8)doi:10.2215/CJN.15040920

2. Randhawa PS, Shapiro R, Jordan ML, Starzl TE, Demetris AJ. The Histopathological Changes Associated with Allograft Rejection and Drug Toxicity in Renal Transplant Recipients Maintained on FK506. The American Journal of Surgical Pathology. 1993;17(1):60–68. doi:10.1097/00000478-199301000-00007

3. Vincenti F, Rostaing L, Grinyo J, et al. Belatacept and Long-Term Outcomes in Kidney Transplantation. New England Journal of Medicine. 2016;374(4):333–343. doi:10.1056/nejmoa1506027

4. Woodle ES, Kaufman DB, Shields AR, et al. Belatacept-based immunosuppression with simultaneous calcineurin inhibitor avoidance and early corticosteroid withdrawal: A prospective, randomized multicenter trial. American Journal of Transplantation. 2020;20(4):1039–1055. doi:10.1111/ajt.15688

5. M. C-RC, Godarova A, Shi T, et al. mTOR Inhibitor Therapy Diminishes Circulating CD8+ CD28-Effector Memory T Cells and Improves Allograft Inflammation in Belatacept-refractory Renal Allograft Rejection. Transplantation. 2020 May 2020;104(5)doi:10.1097/TP.0000000000002917

6. F. V, Charpentier B, Vanrenterghem Y, et al. A phase III study of belatacept-based immunosuppression regimens versus cyclosporine in renal transplant recipients (BENEFIT study). American journal of transplantation : official journal of the American Society of Transplantation and the American Society of Transplant Surgeons. 2010 Mar 2010;10(3)doi:10.1111/j.1600-6143.2009.03005.x

7. Ristov J, Espie P, Ulrich P, et al. Characterization of the in vitro and in vivo properties of CFZ533, a blocking and non-depleting anti-CD40 monoclonal antibody. American Journal of Transplantation. 2018;18(12):2895–2904. doi:10.1111/ajt.14872

8. P. U, Flandre T, Espie P, et al. Nonclinical Safety Assessment of CFZ533, a Fc-Silent Anti-CD40 Antibody, in Cynomolgus Monkeys. Toxicological sciences : an official journal of the Society of Toxicology. 11/01/2018 2018;166(1)doi:10.1093/toxsci/kfy196

9. F. C, Wieczorek G, Audet M, et al. A novel, blocking, Fc-silent anti-CD40 monoclonal antibody prolongs nonhuman primate renal allograft survival in the absence of B cell depletion. American journal of transplantation : official journal of the American Society of Transplantation and the American Society of Transplant Surgeons. 2015 Nov 2015;15(11)doi:10.1111/ajt.13377

10. Mohiuddin MM, Singh AK, Corcoran PC, et al. Role of anti-CD40 antibody-mediated costimulation blockade on non-Gal antibody production and heterotopic cardiac xenograft survival in a GTKO.hCD46Tg pig-to-baboon model. Xenotransplantation. 2014;21(1):35–45. doi:10.1111/xen.12066

11. L. H, Mathews D, Breeden CA, et al. Pre-transplant antibody screening and anti-CD154 costimulation blockade promote long-term xenograft survival in a pig-to-primate kidney transplant model. Xenotransplantation. 2015 May-Jun 2015;22(3)doi:10.1111/xen.12166

12. Mohiuddin MM, Singh AK, Corcoran PC, et al. Chimeric 2C10R4 anti-CD40 antibody therapy is critical for long-term survival of GTKO.hCD46.hTBM pig-to-primate cardiac xenograft. Nature Communications. 2016;7(1):11138. doi:10.1038/ncomms11138

13. Vincenti F, Klintmalm G, Yang H, et al. A randomized, phase 1b study of the pharmacokinetics, pharmacodynamics, safety, and tolerability of bleselumab, a fully human, anti-CD40 monoclonal antibody, in kidney transplantation. American Journal of Transplantation. 2020;20(1):172–180. doi:10.1111/ajt.15560

14. Harland RC, Klintmalm G, Jensik S, et al. Efficacy and safety of bleselumab in kidney transplant recipients: A phase 2, randomized, open-label, noninferiority study. American Journal of Transplantation. 2020;20(1):159–171. doi:10.1111/ajt.15591

15. N. Rp, Plumb TJ, Crowley SD, Coffman TM. Effector mechanisms in transplant rejection. Immunological reviews. 2003 Dec 2003;196doi:10.1046/j.1600-065x.2003.00090.x

16. Morris H, DeWolf S, Robins H, et al. Tracking donor-reactive T cells: Evidence for clonal deletion in tolerant kidney transplant patients. research-article. 2015-01-28 2015;doi:10.1126/scitranslmed.3010760

17. Dewolf S, Grinshpun B, Savage T, et al. Quantifying size and diversity of the human T cell alloresponse. JCI Insight. 2018;3(15)doi:10.1172/jci.insight.121256

18. Zeng G, Huang Y, Huang Y, Lyu Z, Lesniak D, Randhawa P. Antigen-Specificity of T Cell Infiltrates in Biopsies With T Cell–Mediated Rejection and BK Polyomavirus Viremia: Analysis by Next Generation Sequencing. American Journal of Transplantation. 2016;16(11):3131–3138. doi:10.1111/ajt.13911

19. L. L, Wang L, Chen H, et al. T cell repertoire following kidney transplantation revealed by high-throughput sequencing. Transplant immunology. 2016 Nov 2016;39doi:10.1016/j.trim.2016.08.006

20. Alachkar H, Mutonga M, Kato T, et al. Quantitative characterization of T-cell repertoire and biomarkers in kidney transplant rejection. BMC Nephrology. 2016;17(1)doi:10.1186/s12882-016-0395-3

21. Famulski KS, Einecke G, Sis B, et al. Defining the Canonical Form of T-Cell-Mediated Rejection in Human Kidney Transplants. American Journal of Transplantation. 2010;10(4):810–820. doi:10.1111/j.1600-6143.2009.03007.x

22. Reeve J, Sellarés J, Mengel M, et al. Molecular Diagnosis of T Cell-Mediated Rejection in Human Kidney Transplant Biopsies. American Journal of Transplantation. 2013;13(3):645–655. doi:10.1111/ajt.12079

23. F. Hp, Matas A, Kasiske BL, Madill-Thomsen KS, Mackova M, Famulski KS. Molecular phenotype of kidney transplant indication biopsies with inflammation in scarred areas. American journal of transplantation : official journal of the American Society of Transplantation and the American Society of Transplant Surgeons. 2019 May 2019;19(5)doi:10.1111/ajt.15178

24. F. T, Barbacioru C, Wang Y, et al. mRNA-Seq whole-transcriptome analysis of a single cell. Nature methods. 2009 May 2009;6(5)doi:10.1038/nmeth.1315

25. Han A, Glanville J, Hansmann L, Davis MM. Linking T-cell receptor sequence to functional phenotype at the single-cell level. Nature Biotechnology. 2014;32(7):684–692. doi:10.1038/nbt.2938

26. H. W, Malone AF, Donnelly EL, et al. Single-Cell Transcriptomics of a Human Kidney Allograft Biopsy Specimen Defines a Diverse Inflammatory Response. Journal of the American Society of Nephrology : JASN. 2018 Aug 2018;29(8)doi:10.1681/ASN.2018020125

27. F. Ma, Wu H, Fronick C, Fulton R, Gaut JP, Humphreys BD. Harnessing Expressed Single Nucleotide Variation and Single Cell RNA Sequencing To Define Immune Cell Chimerism in the Rejecting Kidney Transplant. Journal of the American Society of Nephrology : JASN. 2020 Sep 2020;31(9)doi:10.1681/ASN.2020030326

28. Snyder ME, Moghbeli K, Bondonese A, et al. Modulation of tissue resident memory T cells by glucocorticoids after acute cellular rejection in lung transplantation. Journal of Experimental Medicine. 2022;219(4)doi:10.1084/jem.20212059

29. Arazi A, Rao DA, Berthier CC, et al. The immune cell landscape in kidneys of patients with lupus nephritis. Nat Immunol. Jul 2019;20(7):902–914. doi:10.1038/s41590-019-0398-x

30. Donlin LT, Rao DA, Wei K, et al. Methods for high-dimensional analysis of cells dissociated from cryopreserved synovial tissue. Arthritis Res Ther. Jul 11 2018;20(1):139. doi:10.1186/s13075-018-1631-y

31. Adam M, Potter AS, Potter SS. Psychrophilic proteases dramatically reduce single cell RNA-seq artifacts: A molecular atlas of kidney development. Development. 2017;144(19):3625–3632. doi:10.1242/dev.151142

32. Stewart BJ, Ferdinand JR, Young MD, et al. Spatiotemporal immune zonation of the human kidney. Science. 2019/09/27 2019;365(6460):1461–1466. doi:10.1126/science.aat5031

33. S. C, Mills J, McGrath MS. Quantitative immunocytofluorographic analysis of CD4 surface antigen expression and HIV infection of human peripheral blood monocyte/macrophages. AIDS research and human retroviruses. 1987 Summer 1987;3(2)doi:10.1089/aid.1987.3.135

34. Behr FM, Beumer-Chuwonpad A, Kragten NAM, Wesselink TH, Stark R, Gisbergen KPJM. Circulating memory CD8 <sup>+</sup> T cells are limited in forming CD103 <sup>+</sup> tissue-resident memory T cells at mucosal sites after reinfection. European Journal of Immunology. 2021;51(1):151–166. doi:10.1002/eji.202048737

35. Kok L, Masopust D, Schumacher TN. The precursors of CD8+ tissue resident memory T cells: from lymphoid organs to infected tissues. Nature Reviews Immunology. 2022;22(5):283–293. doi:10.1038/s41577-021-00590-3

36. Mackay LK, Minnich M, Kragten NAM, et al. Hobit and Blimp1 instruct a universal transcriptional program of tissue residency in lymphocytes. Science. 04/22/2016 2016;352(6284)doi:10.1126/science.aad2035

37. Crowl JT, Heeg M, Ferry A, et al. Tissue-resident memory CD8+ T cells possess unique transcriptional, epigenetic and functional adaptations to different tissue environments. Nature Immunology. 2022;23(7):1121–1131. doi:10.1038/s41590-022-01229-8

38. de Leur K, Dieterich M, Hesselink DA, et al. Characterization of donor and recipient CD8+ tissue-resident memory T cells in transplant nephrectomies. Scientific Reports. 2019;9(1)doi:10.1038/s41598-019-42401-9

39. Khan O, Giles JR, McDonald S, et al. TOX transcriptionally and epigenetically programs CD8+ T cell exhaustion. Nature. 2019;571(7764):211–218. doi:10.1038/s41586-019-1325-x

40. H. Hm, Hoogeboom M, de Paus RA, et al. Redirection of antileukemic reactivity of peripheral T lymphocytes using gene transfer of minor histocompatibility antigen HA-2-specific T-cell receptor complexes expressing a conserved alpha joining region. Blood. 11/15/2003 2003;102(10)doi:10.1182/blood-2003-05-1524

41. C. A, Jelencsics K, Hu K, et al. Prospective Tracking of Donor-Reactive T-Cell Clones in the Circulation and Rejecting Human Kidney Allografts. Frontiers in immunology. 10/14/2021 2021;12doi:10.3389/fimmu.2021.750005

42. Brehm MA, Daniels KA, Priyadharshini B, et al. Allografts stimulate cross-reactive virus-specific memory CD8 T cells with private specificity. Am J Transplant. 2010 Aug 2010;10(8)doi:10.1111/j.1600-6143.2010.03161.x

43. van den Heuvel H, van der Meer-Prins EMW, van Miert PPMC, Zhang X, Anholts JDH, Claas FHJ. Infection with a virus generates a polyclonal immune response with broad alloreactive potential. Hum Immunol. 2019 Feb 2019;80(2)doi:10.1016/j.humimm.2018.10.014

44. Castro-Rojas CM, Godarova A, Shi T, et al. mTOR inhibitor therapy diminishes circulating CD8+ CD28-effector memory T cells and improves allograft inflammation in belatacept-refractory renal allograft rejection. Transplantation. 2020 May 2020;104(5)doi:10.1097/TP.0000000000002917

45. Snyder ME, Finlayson MO, Connors TJ, et al. Generation and persistence of human tissue-resident memory T cells in lung transplantation. Science Immunology. 2019;4(33):eaav5581. doi:10.1126/sciimmunol.aav5581

46. R. Y, El-Asady R, Liu K, Wang D, Drachenberg CB, Hadley GA. Critical role for CD103+CD8+ effectors in promoting tubular injury following allogeneic renal transplantation. Journal of immunology (Baltimore, Md : 1950). 09/01/2005 2005;175(5)doi:10.4049/jimmunol.175.5.2868

47. Satija R, Farrell JA, Gennert D, Schier AF, Regev A. Spatial reconstruction of single-cell gene expression data. Nat Biotechnol. 2015;33(5):495–502. doi:10.1038/nbt.3192

48. Stuart T, Butler A, Hoffman P, et al. Comprehensive integration of single-cell data. Cell. 2019;177(7):1888-1902.e21. doi:10.1016/j.cell.2019.05.031

49. Nazarov V, Tsvetkov VO, Rumynskiy E, Popov AA, Balashov I, Samokhina M. immunarch: bioinformatics analysis of T-cell and B-cell immune repertoires. https://githubcom/immunomind/immunarch. 2022;

50. Kowarz E, Löscher D, Marschalek R. Optimized Sleeping Beauty transposons rapidly generate stable transgenic cell lines - Kowarz - 2015 - Biotechnology Journal - Wiley Online Library. Biotech Method. 2023;doi:10.1002/biot.201400821

