## Supplementary Data for "Single cell transcriptomic analysis of renal allograft rejection reveals novel insights into intragraft TCR clonality"

### Extended Data Figures

Shi *et al.*

A

| Rej. Group | Patient ID | Total No. MM | | Class I HLA | | | | | | No. Class I MM | | Class II HLA | | | | | | No. Class II MM |
|---|---|---|---|---|---|---|---|---|---|---|---|---|---|---|---|---|---|---|
| | | | | A | | B | | C | | | | DRB | | DQB | | DPB | | |
| Rejection under Tacrolimus | TAC_1 | 2 | Patient | A02 | - | B62 | - | C10 | - | 1 | Patient | DR04 | DR11 | DQ07 | DQ08 | DP02:01 | DP04:01 | 1 |
| | | | Donor | A02 | - | **B60** | - | C10 | - | | Donor | DR04 | DR11 | DQ07 | DQ08 | **DP16:01** | DP04:01 | |
| | TAC_2 | 8 | Patient | A02 | A29 | B08 | B45 | C06 | C07 | 3 | Patient | DR04 | DR17 | DQ02 | DQ07 | DP04:01 | DP15:01 | 5 |
| | | | Donor | A02 | - | **B13** | **B62** | C06 | **C10** | | Donor | **DR01** | **DR09** | **DQ05** | **DQ09** | DP04:01 | **DP02** | |
| | TAC_3 | 8 | Patient | A03 | A29 | B44 | B72 | C02 | C16 | 4 | Patient | DR04 | DR07 | DQ02 | DQ07 | DP04:01 | DP10:01 | 4 |
| | | | Donor | **A01** | **A11** | **B35** | - | **C04** | - | | Donor | DR04 | **DR01** | **DQ05** | DQ07 | **DP02:01** | **DP03:01** | |
| | TAC_4 | 8 | Patient | A24 | A32 | B35 | B60 | C04 | C10 | 5 | Patient | DR08 | DR11 | DQ04 | DQ07 | DP03:01 | DP06:01 | 3 |
| | | | Donor | **A02** | - | **B08** | **B50** | **C06** | **C07** | | Donor | DR08 | **DR04** | **DQ08** | DQ07 | **DP04:01** | - | |
| Rejection under Iscalimab | ISCAL_1 | 3 | Patient | A30 | A68 | B53 | B65 | C04 | C08 | 2 | Patient | DR15 | DR17 | DQ02 | DQ06 | DP02:01 | DP17:01 | 1 |
| | | | Donor | A30 | **A32** | **B64** | " | - | " | | Donor | DR15 | **DR07** | DQ02 | DQ06 | DP02:01 | - | |
| | ISCAL_2 | 8 | Patient | A03 | A68 | B07 | B37 | C06 | C07 | 5 | Patient | DR13 | DR15 | DQ06 | DQ06 | DP01:01 | DP04:01 | 3 |
| | | | Donor | A03 | **A02** | **B35** | **B51** | **C04** | **C14** | | Donor | **DR01** | - | **DQ05** | - | **DP15:01** | DP04:01 | |
| | ISCAL_3 | 11 | Patient | A23 | A24 | B14 | B81 | C08 | C08 | 6 | Patient | DR11 | DR12 | DQ05:01 | DQ06:02 | DP105:01 | DP105:01 | 5 |
| | | | Donor | **A01** | **A26** | **B44** | **B57** | **C06** | **C16** | | Donor | **DR07** | - | **DQ02** | **DQ09** | **DP11:01** | **DP13:01** | |
| Rejection under Belatacept | BELA_1 | 6 | Patient | A03 | - | B07 | B62 | C07 | C10 | 3 | Patient | DR01 | DR15 | DQ05 | DQ06 | DP04:01 | DP04:02 | 3 |
| | | | Donor | **A02** | - | **B08** | **B60** | C07 | C10 | | Donor | **DR13** | **DR17** | **DQ02** | DQ06 | DP04:01 | DP04:02 | |
| | BELA_2 | 10 | Patient | A02 | A32 | B60 | B64 | C08 | C10 | 5 | Patient | DR04 | DR13 | DQ06 | DQ07 | DP03:01 | DP04:01 | 5 |
| | | | Donor | **A01** | - | **B08** | **B44** | **C07** | **C16** | | Donor | **DR07** | **DR17** | **DQ02** | - | **DP01:01** | **DP04:02** | |
| | BELA_3 | 10 | Patient | A03 | A30 | B56 | B57 | C01 | C18 | 5 | Patient | DR03:02 | DR10:01 | DQ04 | DQ05 | DP01:01 | DP17:01 | 5 |
| | | | Donor | **A02** | - | **B07** | **B61** | **C02** | **C07** | | Donor | **DR07** | **DR08** | DQ04 | **DQ02** | **DP02:01** | **DP04:01** | |

B

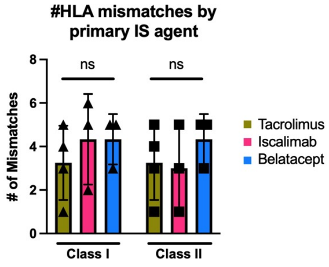


**Extended Data Figure 1. Participant-donor HLA typing.** (A) 2-digit HLA typing is listed for each participant for class I HLAs (left) and class II HLAs (right). Numbers of mismatches range from 2 to 11 total mismatched HLAs. (B) Bar graphs compare total number of class I or class II mismatches between rejection groups (tacrolimus [mustard], iscalimab [pink], belatacept [blue]).

A

| Rej. Group | Patient ID | IS at Time of Biopsy | Biopsy Date (PTD) | Rej. Type | Rej. Grade | Pathology Composite Score | | | | | | | | | | | | | DSA Status (*non-HLA pos) |
|---|---|---|---|---|---|---|---|---|---|---|---|---|---|---|---|---|---|---|---|
| | | | | | | v | t | i | g | ptc | ci | ct | cg | cv | mm | ah | ti | c4d | |
| Rejection under Tacrolimus | TAC_1 | Tac/MMF | 14 | ACR | 1A | 0 | 1 | 2 | 2 | 2 | 1 | 1 | 0 | 1 | 0 | 0 | . | + | Neg |
| | TAC_2 | Tac/MMF/Pred | 1963 | Mixed | 1A | 0 | 2 | 2 | 1 | 2 | 2 | 2 | 1 | 1 | 1 | 2 | 3 | 1+ | Pos(II) |
| | TAC_3 | Tac/MMF/Pred | 217 | ACR | 1B | 0 | 3 | 3 | 1 | 1 | 0 | 0 | 0 | 0 | 0 | 0 | 3 | 1+ | Neg |
| | TAC_4 | Tac/MMF | 809 | ACR | 1B | 0 | 3 | 3 | 0 | 2 | 1 | 1 | 0 | 1 | 1 | 0 | 3 | 0 | Neg |
| Rejection under Iscalimab | ISCAL_1 | Iscal/MMF/Pred | 60 | ACR | 1A | 1 | 3 | 3 | 0 | 1 | 0 | 0 | 0 | . | 0 | 0 | 3 | 1 | Neg |
| | ISCAL_2 | Iscal/MMF/Pred | 14 | ACR | 1A | 0 | 1 | 2 | 0 | 0 | 0 | 0 | 0 | 0 | 0 | 0 | 2 | - | Neg |
| | ISCAL_3 | Iscal/Pred | 137 | ACR | 1B | 0 | 3 | 3 | 0 | 1 | 1 | 1 | 0 | 0 | 0 | 0 | 3 | 1+ | Neg |
| Rejection under Belatacept | BELA_1 | Bela/MMF | 111 | ACR | 2A | 1 | 1 | 1 | 0 | 1 | 1 | 1 | 0 | 0 | 1 | 1 | . | 0 | Neg |
| | BELA_2 | Bela/MMF | 1532 | ACR | 1B | 0 | 3 | 2 | 0 | 1 | 1 | 1 | 0 | 1 | 0 | 0 | 2 | 1+ | Pos(II)* |
| | BELA_3 | Bela/MMF | 172 | ACR | 2A | 1 | 2 | 2 | 0 | 1 | 1 | 1 | 0 | 0 | 0 | 0 | 2 | 0 | Neg |

B

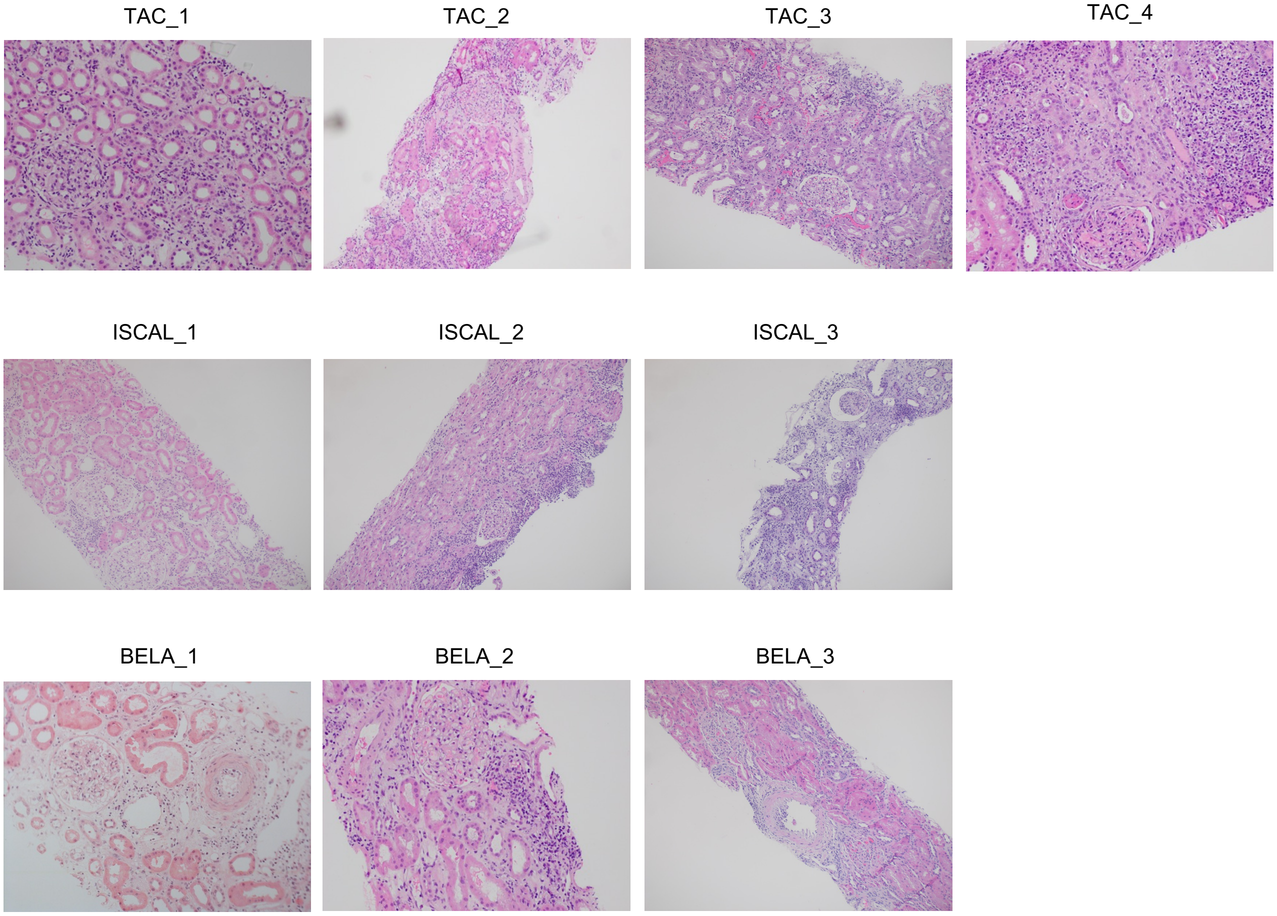


**Extended Data Figure 2. Clinical and pathological characteristics of index rejection biopsies. (A)** Of the 10 index biopsies obtained from participants undergoing rejection, there was an even spread across Banff rejection grades, including Banff 1A, 1B, and 2A rejections. Pathology composite scores and DSA status are provided for each sample. **(B)** Representative histological H&E images of each index biopsy sample reflect their pathological composite scores and Banff rejection grades.

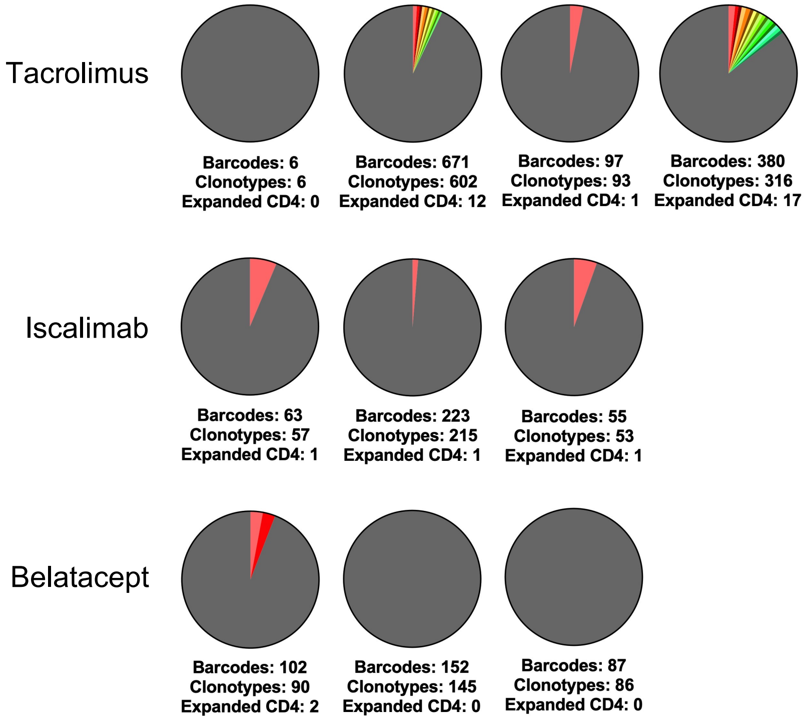


**Extended Data Figure 3. Clonal analysis of CD4+ T cells from index biopsies.** Pie charts display number and frequency of expanded $CD4^+$ clonotypes ($CD4_{EXP}$) found in the biopsy during rejection by participant sample, based on their unique CDR3α/β sequences. Expanded clonotypes are defined as having >2 cells with identical CDR3α/β sequences. Different colors represent individual expanded clonotypes (gray area represents unexpanded clonotypes) and the size of the colored area represents the relative size of the $CD4_{EXP}$.

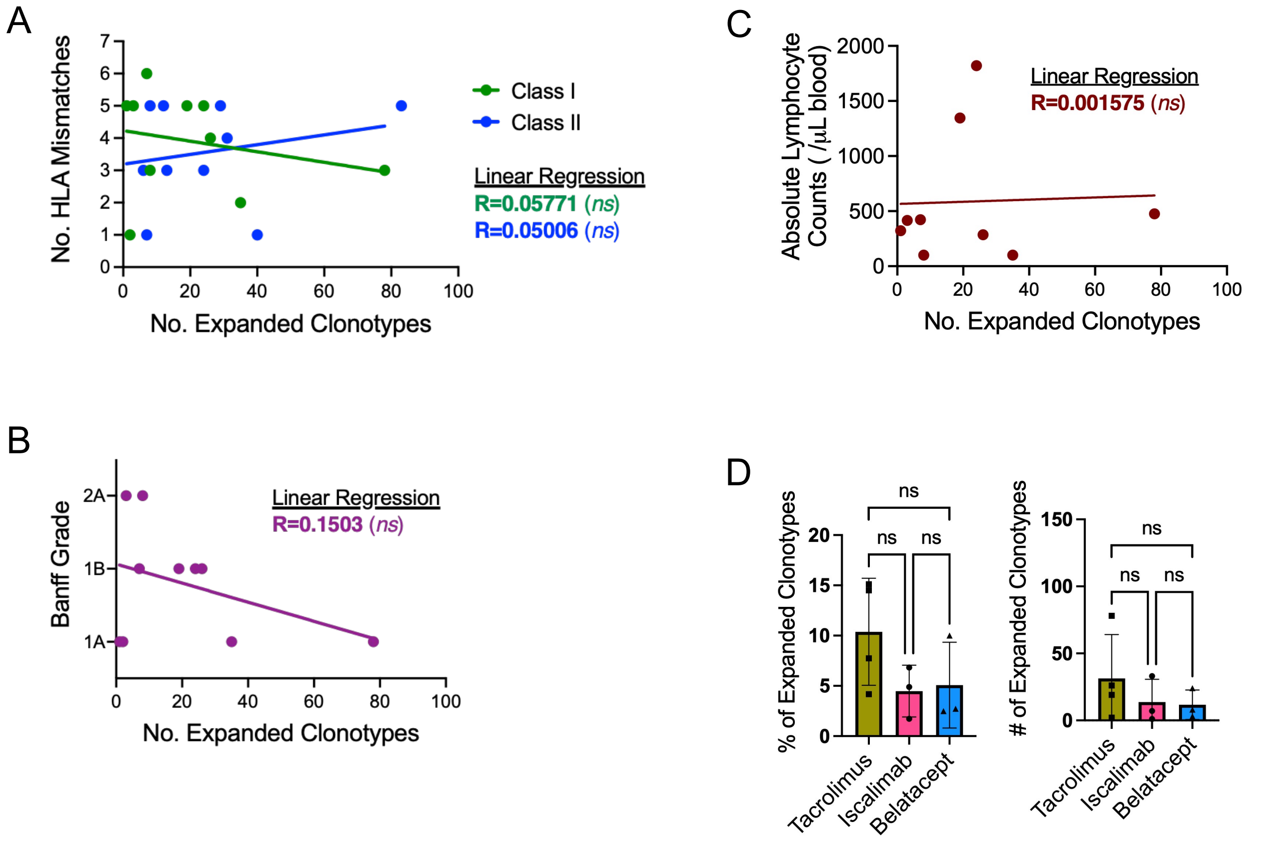


**Extended Data Figure 4. No correlation between numbers of CD8EXP and numbers of HLA mismatches, Banff rejection grade, or post-transplant date of rejection.** (A-C) Simple linear regression analyses were performed on number of CD8EXP against number of (A) HLA mismatches, (B) Banff rejection grade, and (C) post-transplant date (PTD).

| Patient ID | IS at Time of Biopsy | Anti-Rejection Therapy Prior to Biopsy | PTD | Rej Type | ACR Rej Grade | Pathology Composite Score | | | | | | | | | | | | | DSA Status |
|---|---|---|---|---|---|---|---|---|---|---|---|---|---|---|---|---|---|---|---|
| | | | | | | v | t | i | g | ptc | ci | ct | cg | cv | mm | ah | ti | c4d | |
| TAC_3 | Tac/MMF/Pred | . | 217 | ACR | 1B | 0 | 3 | 3 | 1 | 1 | 0 | 0 | 0 | 0 | 0 | 0 | 3 | 1+ | Neg |
| | | rATG; Pred | 232 | ACR | BL | 0 | 1 | 1 | 1 | 1 | 1 | 1 | 0 | 0 | 0 | 0 | 1 | 0 | Neg |
| | | None | 295 | NR | 0 | 0 | 1 | 0 | 0 | 1 | 2 | 2 | 0 | 0 | 0 | 1 | 1 | 1 | Neg |
| ISCAL_1 | Iscal/MMF/Pred | rATG; Pred | 60 | ACR | 1A | 0 | 2 | 1 | 0 | 0 | 0 | 0 | 0 | 0 | 0 | 0 | . | + | Neg |
| | | Tac Added | 78 | NR | 0 | 0 | 1 | 0 | 0 | 0 | 0 | 0 | 0 | 0 | 0 | 0 | . | + | Neg |
| | | Tac Tapered Off | 336 | NR | 0 | 0 | 1 | 0 | 0 | 0 | 0 | 0 | 0 | 2 | 0 | 0 | 0 | 0 | Unk |
| ISCAL_3 | Iscal/Pred | . | 137 | ACR | 1B | 0 | 3 | 3 | 0 | 1 | 1 | 1 | 0 | 0 | 0 | 0 | 3 | 1+ | Neg |
| | | Tac Conversion; Pred | 151 | ACR | 1B | 0 | 3 | 2 | 1 | 0 | 1 | 1 | 0 | 0 | . | 0 | . | 1 | Neg |
| | | MMF Added | 179 | ACR | BL | 0 | 3 | 1 | 1 | 0 | 3 | 3 | 0 | 2 | . | 0 | . | 2 | Neg |
| | | MMF Held | 291 | Mixed | 1B | 0 | 3 | 2 | 1 | 2 | 2 | 2 | 0 | 3 | . | 0 | . | 3 | Pos (I+II) |

**Extended Data Figure 5. Individual participant information for temporal scRNAseq analysis of the response to anti-rejection therapy.** Table displays the type of IS at the time of biopsy, the anti-rejection therapy prior to the biopsy, the day post-transplant (PTD) the biopsy was obtained, the histologic Banff ACR grade including the pathology composite scores, and the DSA status for each participant. Participants TAC_3, ISCAL_1, and ISCAL_3 all received follow-up biopsies following anti-rejection therapy after their index biopsy that revealed ACR. TAC_3 received 2 follow-up biopsies to their initial Banff 1B rejection, ISCAL_1 received 2 follow-up biopsies to their initial Banff 1A rejection, and participant ISCAL_3 received 3 follow-up biopsies to their initial Banff 1B rejection.
